# Strategic Debulking of the Femoral Stem Promotes Load Sharing Through Controlled Flexural Rigidity of the Implant Wall: Optimization of Design by Finite Element Analysis

**DOI:** 10.1101/2024.01.12.575457

**Authors:** Gulshan Sunavala-Dossabhoy, Brent M. Saba, Kevin McCarthy

## Abstract

Hip arthroplasty prostheses are often constructed of metal alloys, and the inherent disparity in the modulus of elasticity between the implant and the femur is attributed to the altered stress-strain pattern in adjacent bone. Rigid implants shield surrounding bone from mechanical loading, and the reduction in skeletal stress required to maintain bone mass and density results in accelerated bone loss, the forerunner to implant loosening and implant failure. Femoral stems of various geometric profiles and surface modifications, materials and material distributions for graded functionality, and porous stem structures have been investigated to achieve mechanical properties of stems that are closer to bone to mitigate stress shielding. For improved load transfer from implant to femur, the proposed study investigated a strategic debulking effort to impart controlled flexibility while retaining sufficient strength and endurance properties of the femoral stem. Using an iterative design process, debulked configurations based on an internal skeletal truss framework were evaluated using finite element analysis as outlined in ISO 7206 standards, with implants offset in natural femur or potted in testing cylinders. The commonality across the debulked designs was the minimization of proximal stress shielding compared to conventional solid implants. Stem topography can influence performance, and the truss implants with and without the calcar collar were evaluated. Load sharing was equally effective irrespective of the collar however, the collar was critical to reducing the stresses in the implant. When bonded directly to bone or cemented in the femur, the truss stem was effective at limiting stress shielding. Nevertheless, a localized increase in principal stress at the lateral proximal junction could negatively affect cement integrity and the bonding of cemented implants. The study determined that superior biomechanical performance of the truss implant is realized with a collared stem that is placed in an interference fit. Mechanistically, the controlled accommodation of deformation of the implant wall provides contextual flexibility and load sharing characteristics to the truss implant.

## 1 Introduction

Prosthetic hip replacement is commonly performed for end-stage hip pathologies including osteoarthritis of the joint, avascular necrosis of the femoral head, and proximal femoral fractures with underlying osteoporosis. Damaged hip joints are replaced with artificial prostheses during hip arthroplasty and the procedure is increasingly performed due in part to osteoarthritis due to a growing aging population and obesity and to the success of the procedure in providing relief from pain and stability to the restored joint (Pivec et al. 2012).

Despite evidence that hip arthroplasty is an excellent treatment option for hip replacement, bone loss around the implants is a concern as mechanical disengagement of the implant occurs, which necessitates revision surgeries (Goldman et al. 2020; Ulrich et al. 2008). Most femoral stems are made of solid metals such as titanium, titanium alloys, cobalt-chrome, or stainless steel that have a modulus of elasticity that is very different from bone (Arabnejad Khanoki and Pasini 2012).

Increased stiffness of the implant results in the bypass of load transfer from the implant to bone and inadequate mechanical loading of bone that is referred to as stress-shielding (Sumner et al. 1998). Stress shielding is the primary cause of reduced peri-implant bone density and the sequelae of bone resorption and implant loosening following hip replacement. The loss of mechanical fixation of the femoral stem that is not due to infection is referred to as aseptic loosening. It is one of the more common shortcomings of hip arthroplasties (Feng, Gu, and Zhou 2022), and revision surgeries are often complicated by the poor quality of residual bone, which increases the risk of intra- and peri- operative fractures. In essence, stress-shielding and associated implant instability reduces the performance and the life expectancy of the prostheses.

The design of the femoral stems and stiffness of the material are important considerations in limiting stress-shielding. Various stem geometries and implant materials have been investigated in an effort to increase load on peri-implant bone and suppress the decline in bone stock. Outer characteristics of the stem such as length (Bieger et al. 2012), taper (Hnat et al. 2009), collar (Jeon et al. 2011), and surface modifications (Heyland et al. 2019) have been assessed for improved bone fixation and osseointegration, but their limited advantages to promoting load transfer have redirected attention to stem materials and material distribution (Arabnejad et al. 2017; Limmahakhun et al. 2017; Wang et al. 2020) to achieve the desired mechanical characteristics. Materials of a low modulus of elasticity improve load transfer, but isoelastic stems with a general increase in flexibility can result in adverse stress concentration in implant and in bone, debonding of the implant from bone, and implant failure (Adam et al. 2002; Trebse et al. 2005). Stems with graded modulus of elasticity either through physicomechanical alteration of material properties or introduction of graded porosities or lattice structures improved load transfer to proximal bone (Limmahakhun et al. 2017; Yamako et al. 2017; Wang et al. 2020). However, achieving the spatial distribution of flexibility and strength without the vulnerability of the design or material, or undetermined time-lapsed changes in bone ingrowth can be a challenge. Based on the premise that stems that accommodate the precise cross-sectional distribution of mechanical sturdiness and flexibility facilitate physiological load bearing, the present study investigated an interior debulking of the stem for improved load sharing in femur.

## 2 Materials and Methods

### 2.1 The Femur and Material Property Derivation

The deidentified CT scans of femurs of a 70-year-old patient were converted to 3D CAD models using 3DSlicer. The model was imported into the Autodesk Meshmixer and mesh tessellation was performed to heal, smoothen, and refine irregularities. Internal smoothening was done using CAD editing features in the Start CCM+ program. The model was imported in Abaqus 2022 HF3 finite element analysis (FEA) program and was converted to a finite element (FE) model. Using Bonemat software, variable density (ρ) and variable modulus of elasticity (E) were mapped from the CT scan data of the femur to the FEA model. Three ρ-E relationships were evaluated and equations (2) and (3) produced similar results.

1. Empirical equation E = 14664 RhoAsh^1.49^ (Morgan, Bayraktar, and Keaveny 2003)
2. Elasticity equation for human femoral neck E = 6850 RhoAsh^1.49^ (Morgan, Bayraktar, and Keaveny 2003)
3. Elasticity equation derived in Javid Dissertation E = 11644 RhoAsh^1.31^ (Javid 2014)

Two representative density values (0.5 and 1.2 g/cm^3^) were taken from the Bonemat exported data file, and corresponding E values with these densities were plotted against the curves of seven E-ρ equations (Javid 2014). Bone mass/density for humans cover a wide range of values, and by validation, equation (3) was found to be the best representative for the current study.

The patient’s CT scan did not include the entire length of the femur, and the distal extension was modeled using visual references from a full human femur (NIH 3D Print Exchange) and was assigned generic bone properties (density = 0.31 g/cm^3^, E = 14700 MPa, v = 0.3).

### 2.2 Femoral Implant Material Properties

Titanium Grade 5 (Ti-6Al-4V) was used for all femoral implant components. Ti-6Al-4V was evaluated for fatigue failure using “fatigue curves”, specific for material, fabrication processes, grain- orientation, testing medium, texture, etc. For Ti-6Al-4V, the worst published texture curve in air was used (Lampman 1996), and based on the fatigue curve, it was determined that at a 10 million cycle run-out, an alternating amplitude stress of 600 MPa is required.

A Keq factor is used for Ti-6Al-4V to account for saline environment testing, manufacturing technique and quality, and other potential surface irregularities. A single “Keq” factor is created from the various contributing “k” factors and is derived from numerous comparisons of the FEA results to laboratory test results for particular products and manufacturing processes, and vendors. A conservative Keq factor of 5 was used in the current study.

### 2.3 FEA Setup

The FEA mesh model of the left femur was cut and reamed to allow for insertion of the femoral implant, with an intervening cement layer (**Figure 1**). The cement region, with its higher quality elements, was changed to bone properties for evaluating direct implant-bone contact. The medium Bonemat properties (Mat-155) were assigned to the adjoining bone (density = 0.764 g/cm^3^, E = 8237 MPa, Poisson ratio (ν) = 0.3). In the models, the femoral head of the implant and the acetabular cap (without the acetabular cap inlay) were included as a single unit.

**Figure 1.**
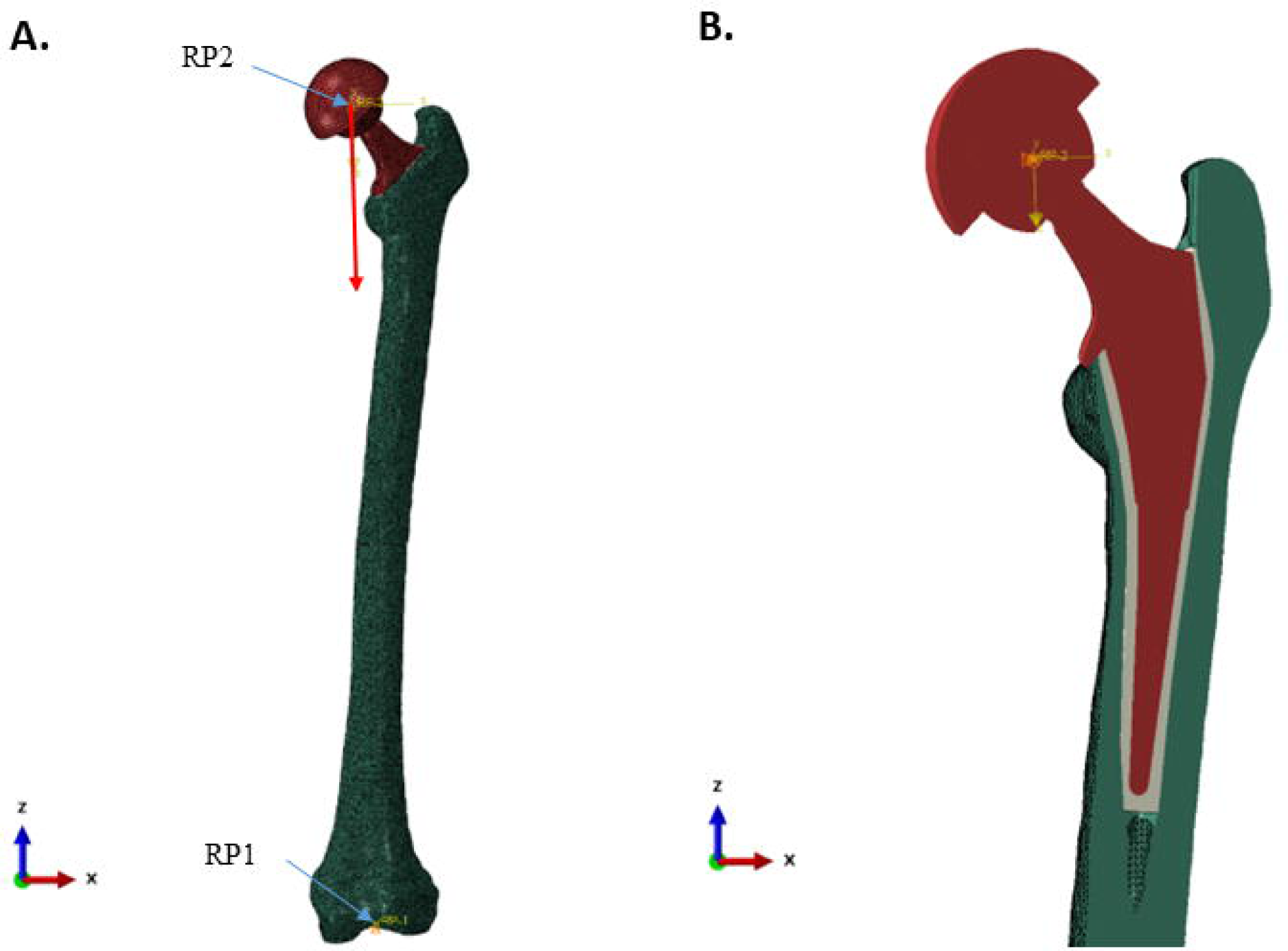
FEA model mesh and materials of implanted femur. A. Smface view of 3D model. Two kinematic couplings were used: RP1, between the condyles of the distal femur and RP2, at the center of the femoral head. RP1 was fixed in space, while displacement of RP2 was constrained to the vertical axis of the local coordinate system *(red an-ow).* B. Vertical section of implanted femur. Tie constraints to bond parts together were used between 1) the caudal and cranial halves of the femur, 2) the acetabular cap and femoral head and neck, and 3) the reamed femur-cement-implant and reamed femur-bone-implant. Smface to smface contact was between the j,nplant collar and mating femur smface.

#### 2.3.1 Meshing, Material Models, Part Tie-Constraints, and Part-to-Part Contact

Quadratic tetrahedral meshing was used for all devices, except for the acetabular cap which was swept linear hexahedral meshed. The femur and the femoral head were meshed with a 1 mm mesh size, while the femoral body and cement were with a 0.75 mm mesh size. All metal components were of Ti-6Al-4V with a linear strain hardening plasticity that had a yield point set to 880 MPa and the ultimate true stress of 1100 MPa and an ultimate true strain of 0.014. Poisson ratio (ν) for all materials was 0.3. Part tie-constraints were used to bond parts together, while surface-to-surface contact was used between the implant collar and the mating femur surface.

#### 2.3.2 Boundary Conditions, Couplings and Spatial Model Constraints

Two kinematic couplings were used in the FEA model. The first coupling (RP1) had the lateral and medial condyles of the distal femur, which were modeled with embedded spheres, controlled by a reference point centered between them. The second coupling (RP2) had the outer surface of the acetabular cap controlled by a reference point located at the center of the femoral head. Loading and displacement-controlled constraints, as assigned to a local Cartesian coordinate section, were applied to the RP2. RP2 movement was limited to the vertical axis of the local coordinate system. RP1 was fixed in space with displacement and rotation of both embedded spheres being restricted (**Figure 1**).

#### 2.3.3 Loads and creation of load direction in the local coordinate system

The loading of the natural femur was accomplished by embedding a small sphere, centered in the femoral head. The load and directional constraint were applied to a reference point at the center of the sphere, controlling its motion. Model loads were applied in accordance with the ISO 7206 Standards, with ISO 7206-4:2010 being most applicable for fatigue conditions and ISO 7206- 10:2018 for static load rating (“ISO 7206-4:2010: Implants for surgery – Partial and total hip joint prostheses – Part 4: Determination of endurance properties and performance of stemmed femoral components. ” 2010; “ISO 7206-6:2013: Implants for surgery – Partial and total hip joint prostheses – Part 6: Endurance properties testing and performance requirements of neck region of stemmed femoral components. ” 2013; “ISO 7206-10:2018: Implants for surgery – Partial and total hip joint prostheses – Part 10: Determination of resistance to static load of modular femoral heads.” 2018). A load of 2300 N was applied to the femoral head, following the vertical direction of the local coordinate system as defined in ISO 7206-4:2010.

The orientation of the implanted femoral stem was as specified in ISO 7206-4:2010 (“ISO 7206- 4:2010: Implants for surgery – Partial and total hip joint prostheses – Part 4: Determination of endurance properties and performance of stemmed femoral components. ” 2010). The α angle, the angle in the frontal plane between the load axis and the stem axis, and the β angle, the angle in the lateral plane perpendicular to the frontal plane between the load axis and the stem axis, were 10° and 9°, respectively. In brief, various axes and planes were used to make the α angle determination in the 3D CAD model using trigonometric relationship. Due to the irregular spatial positioning of the femurs in the original CT data, β angle was measured directly in 3D CAD program as the angle in the lateral plane perpendicular to frontal plane between the load axis and the stem. A new coordinate was established by rotating both α and β by the prescribed amounts to achieve ISO 7206-4:2010 requirements. The ISO 7206-4:2010 load of 2300 N was applied along the local vertical axis for single direction loading. The new coordinates were to simply provide the necessary line of action for application of load in the local coordinate system (**Figure 1)**.

In point to failure experiments, the distal aspect of the implant stem was potted in a test cylinder using Delrin (density = 1.41 g/cm3, E = 3100 MPa, and ν = 0.35) as the potting material. Loading was performed as described in ISO 7206-4:2010, but in a displacement-controlled manner until failure. The acetabular cap was forced to follow the vertical direction in the local coordinate system for an arbitrary 50 mm, and a load-*vs*-displacement chart was created, and the failure load determined. Implant endurance load testing was performed similarly in a potted test cylinder, but instead of a static load, a cyclic load of 2300 N was applied, and implant movement restricted to the vertical direction of the local coordinate system.

#### 2.3.4. Solid, hollow, and truss implant model descriptions

External dimensions of the Zimmer M/L Taper Hip Prosthesis were used for the femoral stem and adapted, or not, with a calcar collar. The solid implant version was followed by the creation of a hollowed-out implant with wall thickness of 2 mm. An internal structure was created with trusses that braced the central spine. The central spine was disconnected from the distal aspect of the stem.

Modifications to the spine and trusses included changes in shape and thickness. The distal part of the femoral stem was solid across all implant configurations (**Figure 2**).

**Figure 2:**
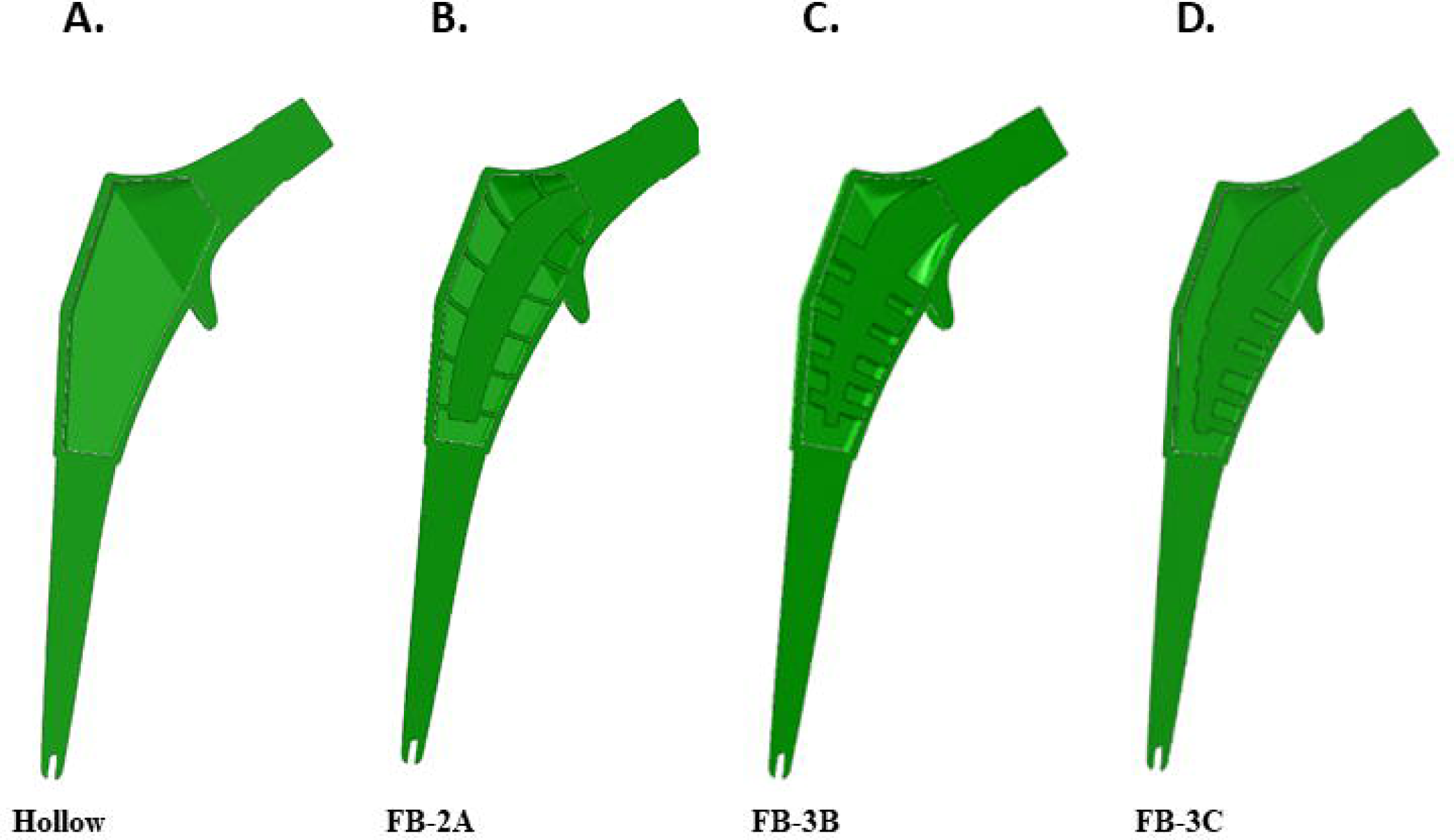
Coronal sectional views of implants. External dimensions were adapted from the Zimmer *NVL* Taper Hip Prosthesis with or without the addition of a collar. Various truss implant designs were created, and sectional views of collared implants are shown. **A.** Proximal hollowed stem. **B-D.** Truss designs, FB- 2A, FB-3B, and FB-3C. The distal portion of all stems was solid

## 3 Results

### 3.1 Stresses in the intact femur

The general distribution of stress after unidirectional loading of the natural femur resulted in higher compressive stress at the medial side of the femur, above the lesser trochanter which extended distally to the diaphysis (**Figure 3A)**. A noticeable increase in tensile stress was observed at the lateral side of diaphysis and at the superior aspect of the neck where it unites with the greater trochanter (**Figure 3B**).

**Figure 3.**
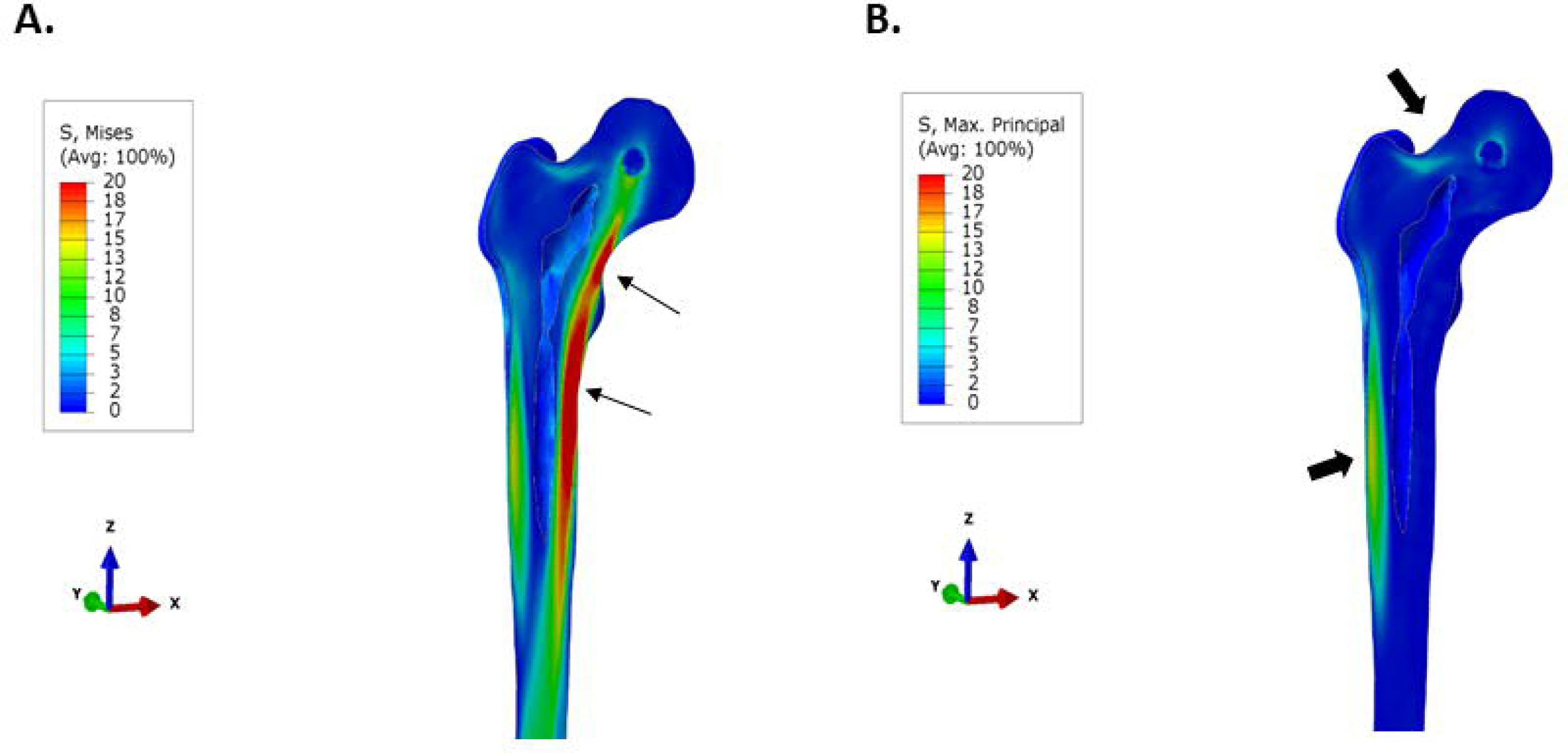
Stress **distribution iu natural femur.** A small sphere was embedded at the center of the femoral head and the load and directional constraints were applied at the center of the sphere. As defined in ISO 7206:4:2010, a load of 2300 N was applied to the femoral head. **A.** Von Mises stress intensity distnbution. Regions of higher compressive stress occur on medial aspect of the femur, above and below the lesser trochanter *(arrows).* **B.** Maximum Principal (tensile) stress plot. The largest region of tensile stress occurs along the lateral aspect of the diaphysis, while it is evident also at the junction of the femoral neck and the greater trochanter *(block anows).* The stress plots are set between O and 20 MPa for comparative purposes.

### 3.2 Stresses in the implanted femur

FEA simulations were conducted for the solid, hollow, and different truss implants under identical setup and loads. Two sets of simulations were conducted for each implanted femur: first, with direct implant-bone bonding and second, with a cement layer between the reamed bony canal and the implant.

#### 3.2.1 Uncemented *vs.* Cemented Stems

In the femur that was contiguous with the collared implant, there was negligible stress in proximal bone with the solid design (**Figure 4A**). The benefit of implant debulking in mitigating stress shielding was clear. Debulking, in general, returned stresses to the region (**Figure 4B-E**). In implants with thicker trusses, FB-3B and FB-3C, stress extended to the anterior and posterior sides of the proximal femur compared to the thin truss implant, FB-2A (**Figure 4**). There was a modest increase in tensile stress localized to the lateral-proximal junction with the hollow and truss implants as compared to the solid implant (**Figure 5**). However, the hollow design had a higher maximum principal stress than any truss design, and the peak tensile stress, 26 MPa, was the same for all truss implants (**Figure 5C-E**).

**Figure 4.**
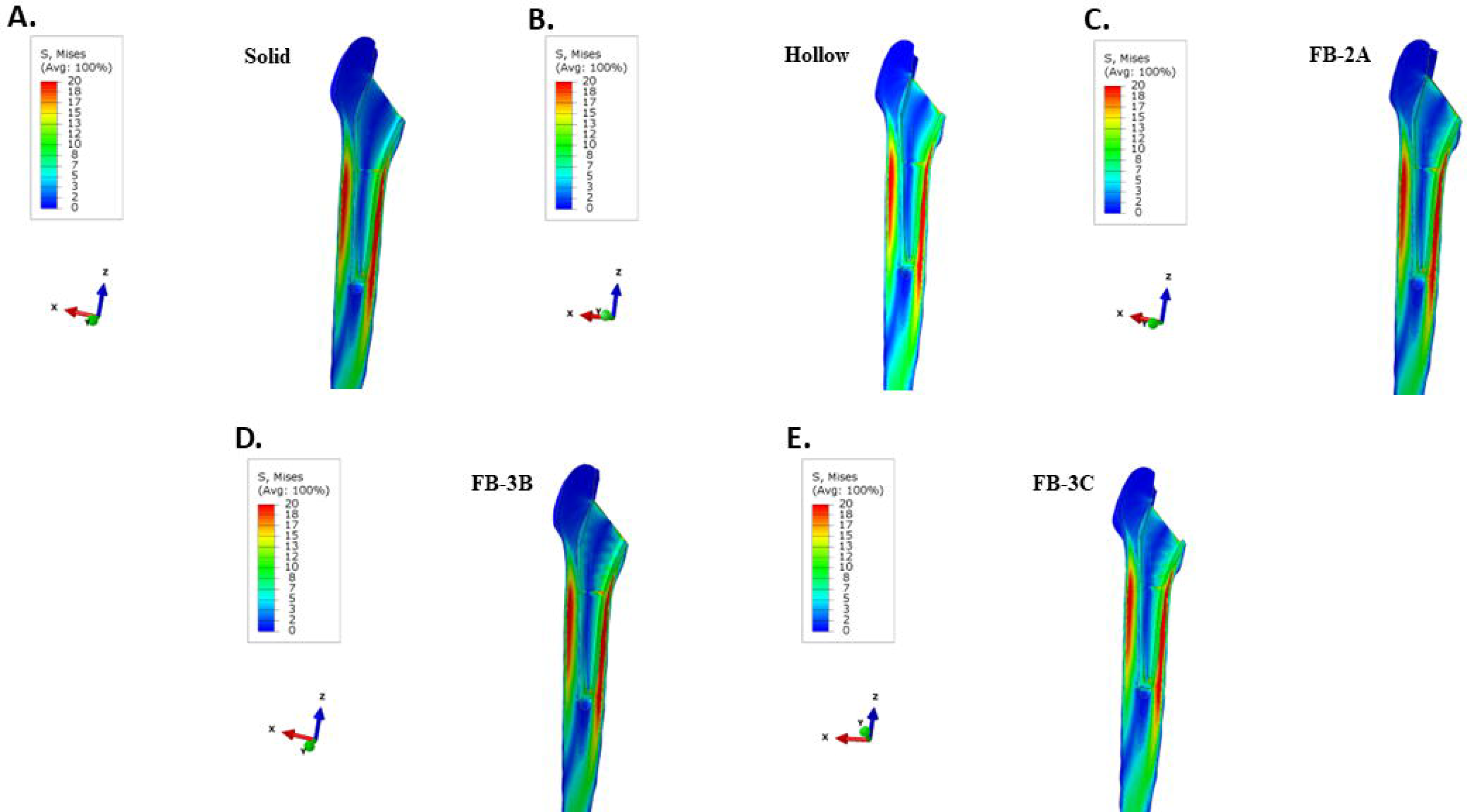
Von fises stress intensity in implanted femur ” th unc.ewented c.ollared stews.A. Solid, B. hollow, C. FB-2A, D. FB-3B, E. FB-3C.The van Mises stress intensity and the maximum (tensile) principal stress plots are set between 0-20 MPa for comparative purposes.

**Figure 5.**
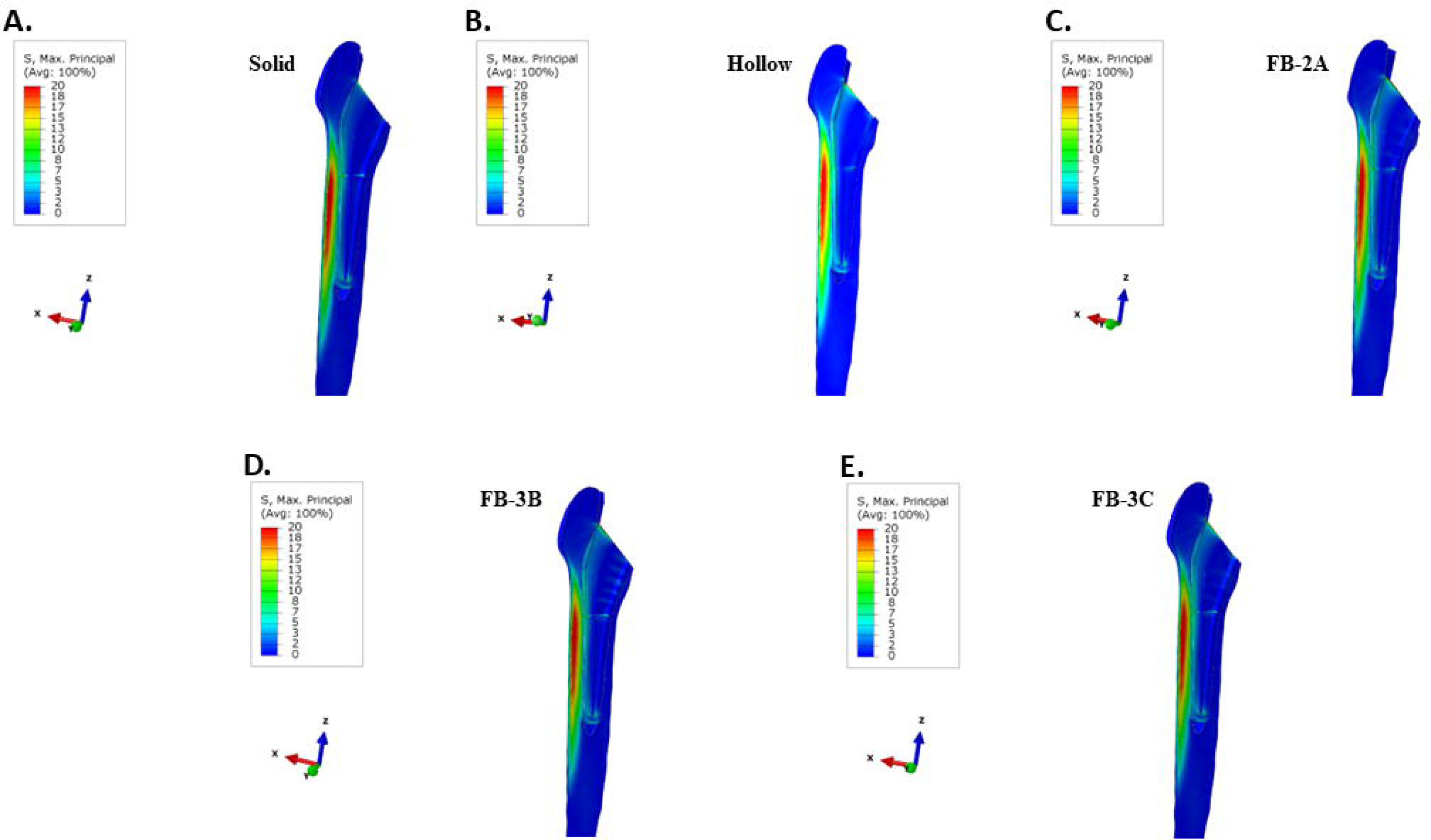
Maximum principal (Tensile) stresses in femur ” th uncemented collared stems.A. Solid, B. hollow, C. FB-2A, **D.** FB-3B, E. FB-3C.All stress plots are set between 0-20 l’vIPa for comparative pwposes.

Cement has a low modulus of elasticity, and it can redistribute the load to the femur. Not surprisingly, mitigation of stress-shielding was significant across all implants when cemented in femur (**Supplementary figure 1**). There was however greater load sharing in proximal region with hollow and truss implants compared to the solid form. The tensile stress profiles were similar across various designs with the highest tensile stresses at the lateral diaphysis (**Supplementary figure 2**). Of note was the occurrence of a highly localized area of tensile stress at the lateral-proximal connection between the cement and the femur when hollow or truss designs were implanted (**Supplementary figure 2B-E**). The hollow design resulted in a higher tensile stress in this region, and differences in flexural rigidity with the truss designs showed an incremental decrease in stress at the location.

The tensile strength for bone-cement bonding is less relative to the bonding between Ti-6Al-4V and cement. The mean tensile strength for bone-cement bonding was calculated as 6.3 MPa, with a minimum value of 2.4 MPa (Beckmann et al. 2014). The cement interface (the surface bonded to bone in the FEA model) was therefore isolated and examined for maximum principal stress, and stress plotted on a scale of 0-2.4 MPa -- 2.4 MPa being the lower value for cement-bone tensile strength (Beckmann et al. 2014). Results showed that the stiffer the implant, the lower the maximum tensile strength at cement-bone interface: 4.1 MPa for the solid, 4.4 MPa for the stiffest truss implant (FB-3B), 4.5 MPa for the remaining truss implants (FB-2A and FB-3C), and 5.3 MPa for the hollow design (**Supplementary figure 3**). A significant increase in tensile stress above the minimum debonding strength of 2.4 MPa occurred with all implant types. Unlike the solid implant, the peak tensile stresses for the hollow implant and truss implants were located at the proximal lateral junctions (**Supplementary figure 3B-E)**. Cement has a low bond strength and can be readily damaged with implants of lower stiffness. The higher tensile stress can affect cement-bone bond and the stresses at the lateral-proximal junctions with debulked designs could de-bond the cement from the femur.

#### 3.2.2 Collar *vs.* Collarless Femoral Stems

To examine the impact of the calcar collar on stress distribution in bone, the solid and truss implants without the calcar collar were investigated. Despite the absence of a calcar collar, FB-3C truss implant when placed in direct interference fit showed improved load sharing in proximal bone compared to the solid implant (**Figure 6A-B**), while maximum principal stress between the collared and collarless solid and truss implants was near-comparable (**Figure 5 and Figure 6C-D**). Alternatively, no difference in Von Mises stress intensity or maximum principal stress was observed with collarless solid or FB-3C stems that were cemented (**Supplementary** Figure 5). There was, however, an increase in cement-bone bonding stress with the collarless stems compared to collared stems (solid collarless: 4.3 MPa *vs.* solid collared: 4.1 MPa; FB-3C collarless: 4.8 MPa *vs.* FB-3C collared:4.5 MPa) (**Figure 3 and Supplementary** Figure 6A**, B**). The higher stresses in the cement layer with collarless stems can disrupt bonding to bone and therefore considered less desirable.

**Figure 6.**
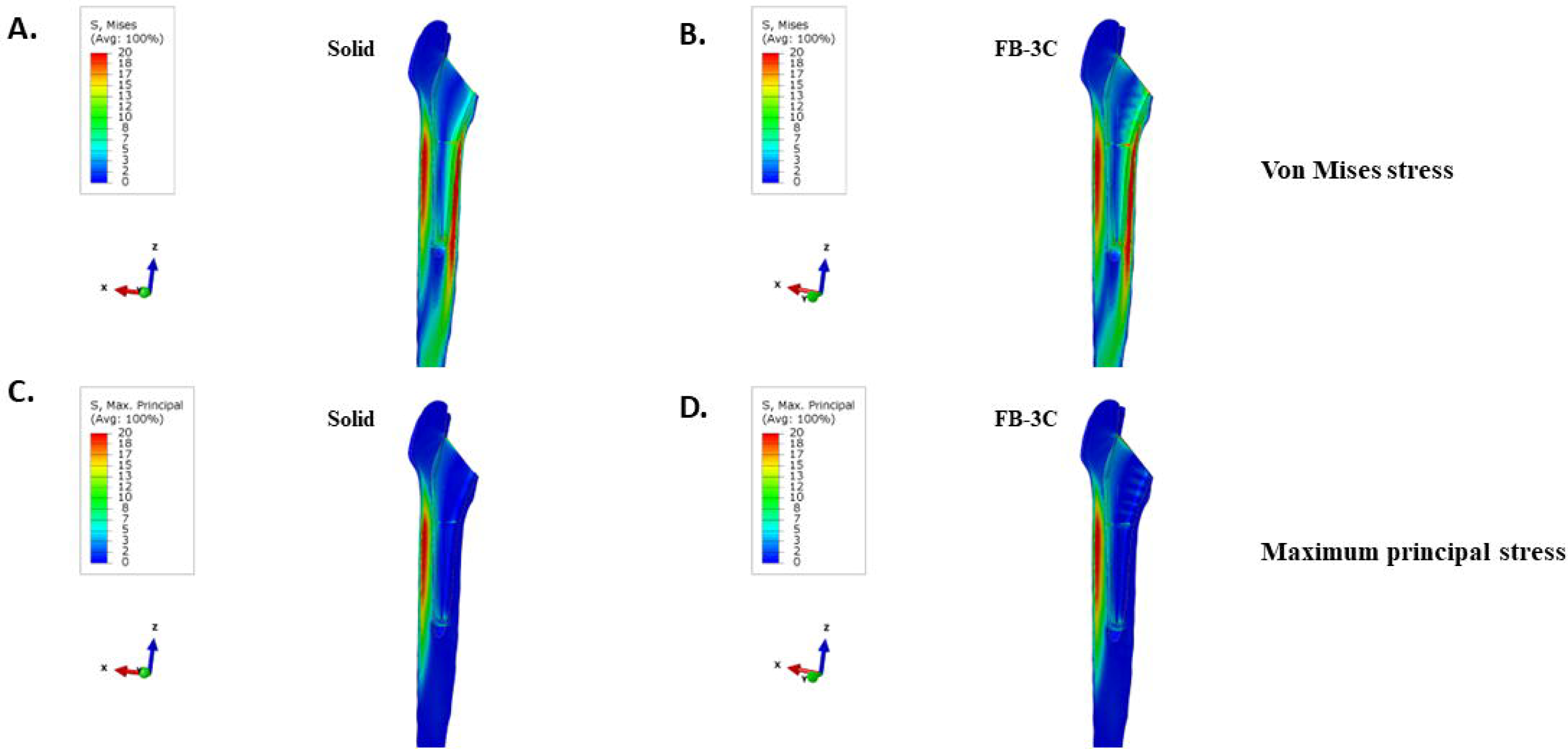
Collarless femoral stems in bone-implant interference fit. A, B. Von Mises stresses and C, **D.**maximum principal stresses in bone. A, C. Solid implant, B, D. FB-3Cimplant. All stress plots are set between 0-20MPafor comparative purposes.

### 3.2 Maximum Principal (Tensile) Stresses in the Implant

To screen for fatigue of the implant, maximum principal stresses in the implant were analyzed. Irrespective of whether the collared implants were directly bonded to bone or cemented, peak stresses at the implant neck were lowest in the most flexible, hollow design, 122 MPa (**Figure 7 and Supplementary** Figure 4). The solid implant showed the highest peak stress in uncemented and cemented conditions, 182 MPa and 184 MPa, respectively, although the area of peak stress was at the implant neck in interference fit while at the lateral commissure between distal and proximal halves in cemented. The truss designs with variations in flexural stiffness had peak stresses between the extremes of the solid and hollow implants. Unlike peak tensile stress at the condylar head-neck junction in solid and hollow implants, the peak stress in truss designs was invariably at the most proximal aspect of the central spine. Among the truss designs, FB-3C exhibited the lowest peak tensile stress -- 158 MPa when cemented and 159 MPa when directly bonded to bone (**Figure 7 and Supplementary** Figure 4). As lower stress in implant can extend implant life, FB-3C was considered the most-favorable truss design.

**Figure 7:**
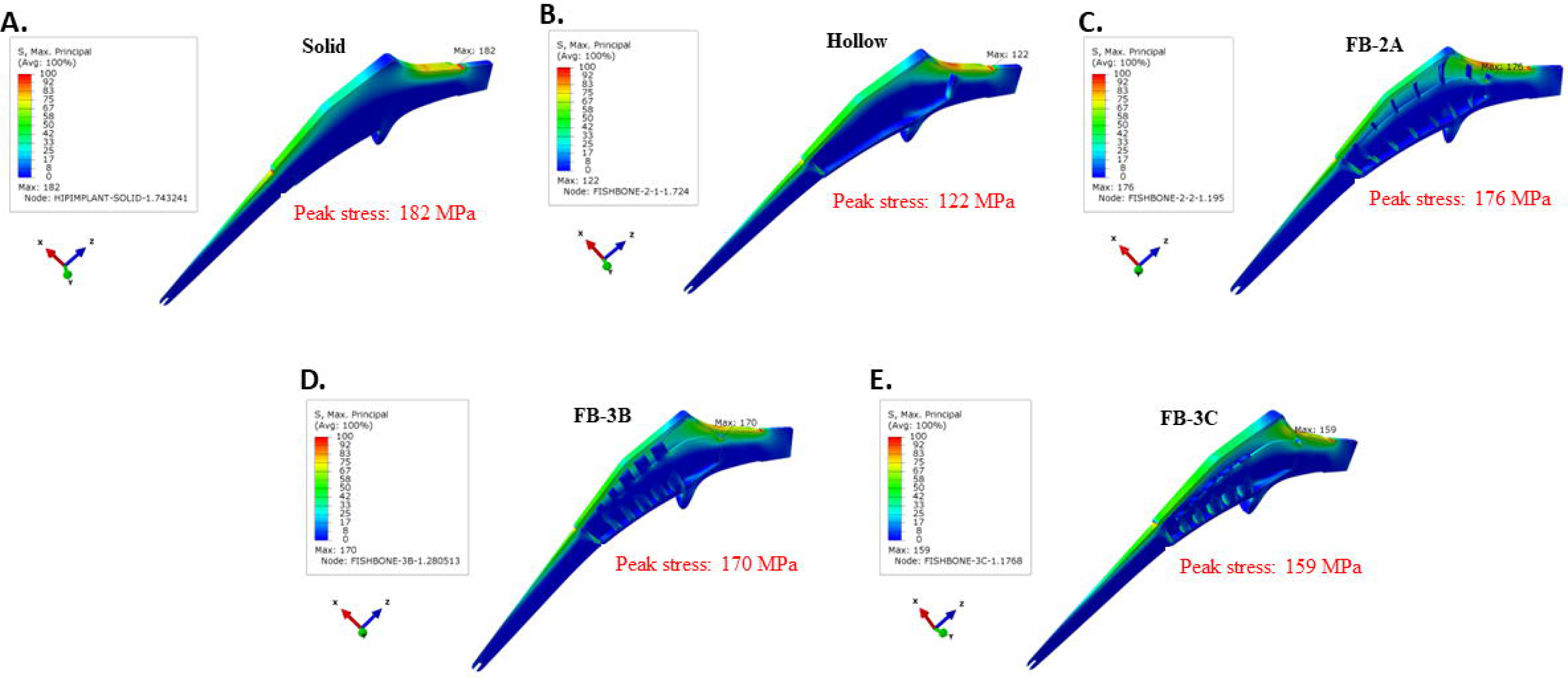
Maximum principal stresses in uncemeoted collared Ti-6Al-4V implants. **A.** Solid, **B.** hollow, C. FB-2A, **D.**FB-3B, **E.** FB-3C. All stress plots are set between 0- 100 MPafor comparative purposes. All stress plots are set between 0-100 MPafor comparative purposes.

Importantly, absence of a collar resulted in increased implant stress in FB-3C. Compared to the peak tensile stress of the collared stem (159 MPa), stress significantly increased in the collarless state (194 MPa) (**Figure 7 and Supplementary** Figure 6D). The maximum principal stress in collarless FB-3C was higher than collared and collarless solid implants (194 MPa *vs.* 182 MPa and 183 MPa, respectively). The results indicate that the calcar collar is critical to the performance and life of the truss implant.

### 3.3 Quantification of Stress in Proximal Bone

Stresses in lateral proximal bone and medial proximal bone along the reamed bone cavity were evaluated with implants in interference fit. Node values along the medial and lateral internal paths were averaged for every 5 mm segment in the cranial-caudal direction. As anticipated, the peri-implant proximal bone was under compressive stress on the medial side, whereas under tensile stress on the lateral side. The compressive stress in medial bone across all segments in Gruen zone 7 was significantly increased with the truss implants compared to the solid configuration irrespective of the stem being collared and collarless (**Figure 8B**). The maximal principal stress (tensile) in lateral proximal bone, Gruen zone 1, was increased with the truss implants, with the increase more noticeable in the most proximal segment (**Figure 8C**). Unlike differences between the solid and FB- 3C, the presence of a calcar collar had no significant impact on compressive or tensile stress distribution in bone. The FB-3B and FB-3C stem designs with calcar collars showed comparable stress distribution in proximal medial and lateral bone (data not shown).

**Figure 8.**
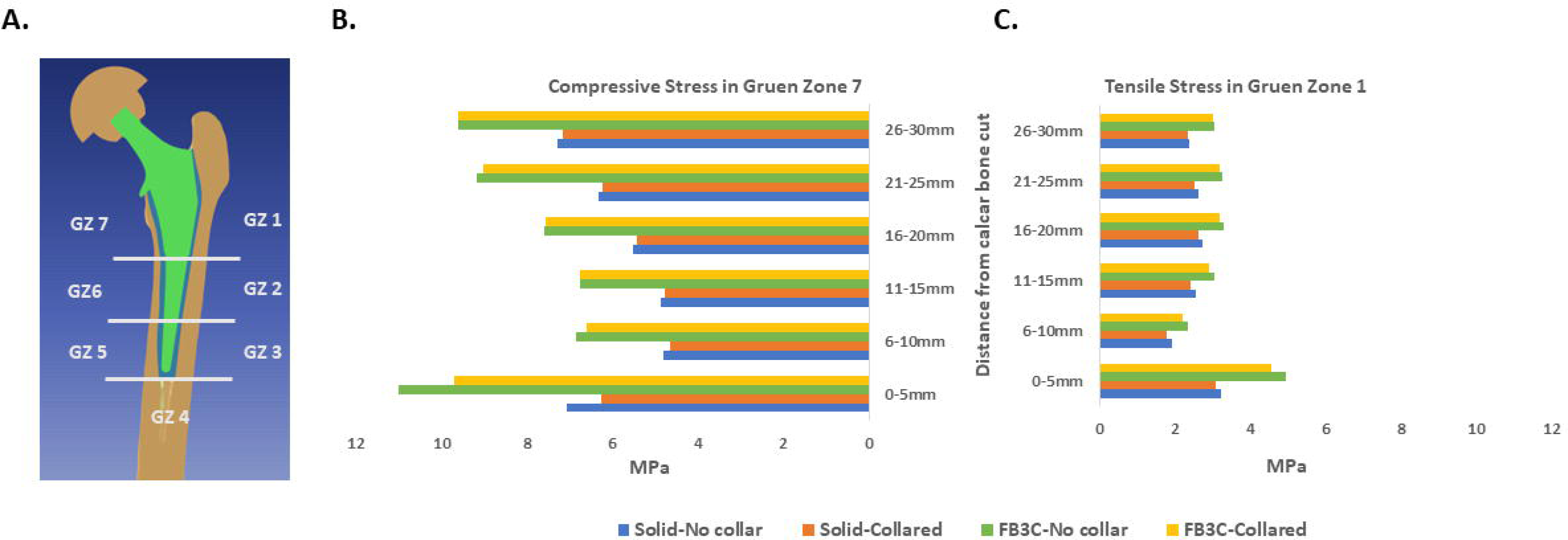
Stress intensity in proximal bone with collared and collarless implants of solid and truss configuration. A. The Gruen reference zones. Nodal stress values were averaged across 5 mm segments in cranial-caudal direction of the reamed bone cavity. B. Minimum principal stress (compressive) in medial bone (Gruen zone 7), C. Maximum principal stress (tensile) in proximal lateral bone (Gruen zone 1).

The medial-caudal flexure of the condylar neck increases compressive stresses on the medial side of the femoral neck and the diaphysis, while it increases tensile stresses at the femoral neck adjoining the greater trochanter (**Figure 3B**). The bio-mimic flexure of the truss implant creates stress patterns similar to natural femur, and the tensile stress in order of 5 MPa generated at the proximal lateral junction is well within the tensile yield strength of bone (51-66 MPa). The load sharing features of truss design are considered beneficial to bone integrity and implant fixation.

### 3.4 Implant Static Load Tests to Failure

The distal ends of the solid implant and the truss implants were potted in the test cylinder and loaded similarly to previous ISO 7206-4:2010 tests, but herein, in a displacement-controlled manner until failure. The conjoined unit of the acetabular cap and the implant head were forced in the vertical direction in the local coordinate system for 50 mm. The solid implant failure occurred at 85 kN due to neck gross yielding, while truss implants, FB-3B and FB-3C, failed at 63 kN due to local buckling of medial implant wall near the potting level (**Figure 9A-B**). Based on the plastic creep analysis, both implants were deemed suitably strong as the failure load exceeded the 2300 N test load in ISO standard 7206-4:2010 by >25 fold.

**Figure 9.**
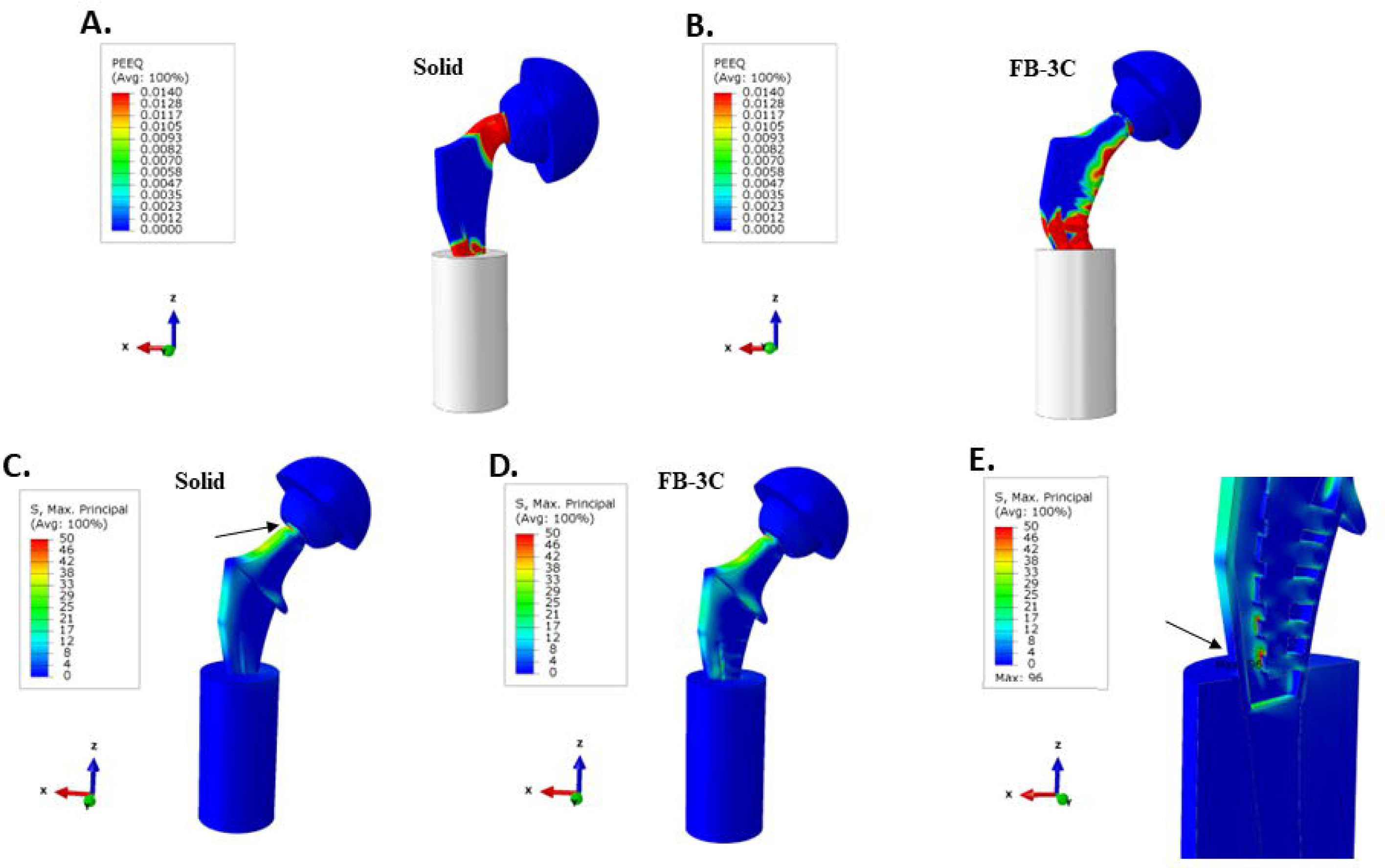
Fatigue tests as per ISO standard set-up. A-B. Static load tests of solid implant and FB-3Cimplant. C-D. Maximum principal stresses in endurance tests of solid implant and FB-3C implant. E. Expanded view of coronal section of FB-3C implant. *Arrows* indicate regions of peak stresses.

### 3.5 Implant Endurance Load Tests

The implant wall, central spine, and trusses of FB-3C were joined together in 3D CAD with fillets added through the inner implant cavity. The stresses are mesh sensitive and based on earlier experience, a mesh size of 0.05 to 0.10 mm at critical locations in fatigue models was used to provide acceptable peak stress results. On cyclic loading of 2300 N, the solid implant displayed peak tensile stress of 65 MPa at the superior surface of the condylar head-to-neck transition (**Figure 9C**). As the loading was unidirectional (not reverse bending), the alternating stress amplitude was computed as 33 MPa. Despite a relatively high-end single K-factor of 3.0, the stress of ∼100 MPa was determined to be well below the limit of alternating stress amplitude of 600 MPa to reach 10 million cycles.

Due to the increased flexibility of FB-3C truss implant, the transition region at the neck had lower peak stress (**Figure 9D**). As the loading was strongly focused on the lower distal portion of the implant that was immobilized, the lower trusses of the skeletal framework showed a higher peak stress of 96 MPa (**Figure 9E**). This stress was not present in previous tests in implants placed in the femur as the loading set-up was different. When embedded in the femur, much of the load was shouldered by the proximal abutting bone. Being potted low on the distal stem for ISO endurance testing, a greater load was placed on the part of the implant not expected to bear such load especially, with implants that support proximal bone integrity through load sharing. Nonetheless, endurance calculations using a highly conservative single K factor of 5.0 to account for unknowns in fabrication technique, an alternating stress amplitude of 240 MPa was calculated. With a 10 million cycle endurance limit of 600 MPa, the stress state of the truss implant was considered low enough to achieve the desired 10 million cycles of loading.

## 4 Discussion

Hip arthroplasties have had a high overall success rate as they effectively reverse joint disability. Rates of primary hip replacements have continued to increase but so have the revision surgeries. Hip revision surgeries are commonly performed because of loss of implant fixation in the bone (“American Joint Replacement Registry (AJRR): 2022 Annual Report. ” 2022; Deere et al. 2022).

Often revision surgeries are complex, and the outcomes are poorer than that for primary hip replacements (Deere et al. 2022). Rigid implants have a propensity for stress shielding, and without a perfect fit between the implant and the reamed femoral canal, rigid stems generate hard and light contact zones which promote unequal loading of bone and osteolysis over time. An implant that is flexible in contrast has the benefit of an improved interference fit that promotes consistent contact loading and a more uniform distribution of stresses. Although the effect of interference fit was not tested in the current work, it is surmised that a reduction in stem stiffness generated either through alteration of implant material properties or a debulking effort will create a compressive stress field at the bone interface, reducing the detrimental effects of tensile stresses in bone.

Various materials of lower Young’s modulus, variations in anatomic surface features, hollowed stems, and stems with regular, graded, or stochastic porosities and lattice designs have been investigated to decrease rigidity and overcome the paucity in stress sharing associated with conventional stems (Arabnejad et al. 2017; Gross and Abel 2001; Heyland et al. 2019; Limmahakhun et al. 2017; Yamako et al. 2017; Yang et al. 2009; Liu et al. 2021). As peri-implant bone loss occurs mostly in the proximal femur, the femoral stem in the study was developed with an internal truss design restricted to the proximal stem, while the distal end was solid. An external implant wall encased the internal structure and provided a continuous contact surface with bone. Our results suggest that through controlled accommodation of deformation of the implant wall, the design promotes load sharing in femur.

To counter the weakness of the hollow design while minimizing stress shielding in proximal bone, the truss design was modeled, and after testing several iterations, FB-3B and FB-3C implants provided the best combination of structural rigidity, selective flexibility, and load sharing characteristics. Although both designs had maximum principal stresses in the implant that were lower than the solid implant, the hemi-truss frame of FB-3C resulted in the lowest implant stress. It was therefore examined further to assess the impact of the calcar collar and cement fixation on stress distribution in bone. However, the balanced truss framework of FB-3B could offer stronger mechanical strength against torsional loads, and the higher implant stresses could be designed out in the model.

It is expected that implants that simulate the mechanical behavior of the natural femur will create compressive stresses in medial proximal bone and in corollary, tensile stresses at the superior surface of the femoral neck and lateral aspect of the greater trochanter. The collarless truss implant was efficient in limiting stress shielding, but the increase in implant stress suggested the necessity of a calcar collar. An issue with cement bonding is material deterioration, and the poor tensile strength of cement increases the propensity of early development of cracks in the cement and the premature loss of implant stability with load-adaptive implant designs. Femoral stems with decreased rigidity as in the case of the truss implant would perform better and have extended usable life when placed in an inference fit. The two salient observations that emerged from the study were that 1) the added flexibility of the truss implant precludes the use of cement for bonding, and 2) the calcar collar decreases implant stresses, a necessity for a longer implant life. For optimal functionality, the truss femoral design benefits from a collared stem that is press fitted in bone.

Biomechanical study of stress distribution identified a major reduction in proximal cortical strain in implanted femurs (Bieger et al. 2012). The reduction in strain is indicative of poor loading of bone. It is therefore not surprising that following hip arthroplasty, the most significant loss of bone mineral density irrespective of femoral stem geometries occurs in proximal periprosthetic bone. Bone loss was most pronounced in medial proximal bone, Gruen zone 7, while the degree of bone loss was less distally, in Gruen zones 3-5 (Inaba et al. 2016). These observations are in alignment with our study where considerable stress shielding was observed on proximal-medial side with the completely solid implant. The restoration of bone loading with the new stem design could positively influence bone integrity and implant lifespan.

The static strength of any debulked material will be inherently weaker than a solid configuration of the same cross-sectional design. The fatigue study as per ISO standards simulates the worst-case scenario in which the implant support is limited to the distal third of the stem. It however does not truly replicate a situation where a bio-mimic implant mitigates stress shielding and in turn, preserves proximal bone integrity. Nevertheless, the *in-silico* analysis suggests that the truss implant is of suitable strength for the expected loads predicted to act on the body. Importantly, improved interference fit of the truss implant in bone would allow for immediate load sharing. Early rehabilitation regimens after uncemented hip replacement favored delayed full weight-bearing to encourage bone growth and stem integration. Delayed weight bearing can accentuate bone deterioration, and a greater degree of post-operative bone loss, at three months post-operatively, was found when partial weight bearing was recommended (Boden and Adolphson 2004). Although subsequent clinical trials have showed no discernable percent change in bone mineral density irrespective the regimen (Wolf et al. 2010, 2013), a consensus is lacking regarding the regimen to adopt. As mineralization of bone is dependent on mechanical loading, early loading of implant could be beneficial to preserving bone mineral density.

Additive manufacturing provides an opportunity to construct structures of complex geometries that subtractive manufacturing is unable to. The construction of 3D lattices and truss designs in metal for medical applications are feasible with the advent of additive fabrication. Build orientation, microstructure, and post-build treatment and surface finish can affect material behavior. The two major factors that lower the performance and life of additively manufactured metal structures are surface roughness and subsurface pores (Afroz et al. 2022; Segersall et al. 2021). Surface-connected porosities are often points of crack initiation and cracks that propagate from the surface into the material can lead to fatal structural failure. In general, the effect of surface roughness and internal defects is more critical for thinner structures, as it can be severely life-limiting. For additively fabricated implants, thin structures can lower fatigue life and be limited in their load bearing capacity. The proposed femoral stem overcomes the limitations of porous designs where additive manufacturing defects can significantly compromise the structural strength of thinner structures. Although completely porous implants have the inherent advantage of bone ingrowth that can add to implant anchorage, the extent of bone invagination and calcification can change the mechanical situation and the predicted performance of the implant. In this regard, an internal truss framework within the femoral stem that is compartmentalized from external influences has the advantage of predictable and consistent performance. It is anticipated that the compressive stresses achieved through interference fit will confer initial implant stability and faster recovery to weight bearing activity, while mechanical deformation of the implant wall will stimulate osteoblastic differentiation of mesenchymal stem cell that is vital to osseointegration and long-term stability of the implant.

## 6 Author Contributions

GSD: conceptualization, created and refined 3D CAD models of femurs, directed the research, and wrote and edited the paper. BMS: created the 3D CAD models of hip implants, generated FE models of femurs and implants, conducted FEA, and wrote the paper. KJM: conceptualization, discussions, and reviewed the paper.

## Acknowledgments

The preliminary work on the hip implant designs was conducted by Behram Dossabhoy, and preliminary models of the solid implant and truss implant were constructed in SolidWorks by Lamont Lackmann (Bossier Parish Community College).

## 8 Supplementary Material

Uploaded separately.

## 9 Data Availability Statement

Not applicable.

