## Supplementary figures and images for "Strategic Debulking of the Femoral Stem Promotes Load Sharing Through Controlled Flexural Rigidity of the Implant Wall: Optimization of Design by Finite Element Analysis"

### Supplemental figure 1

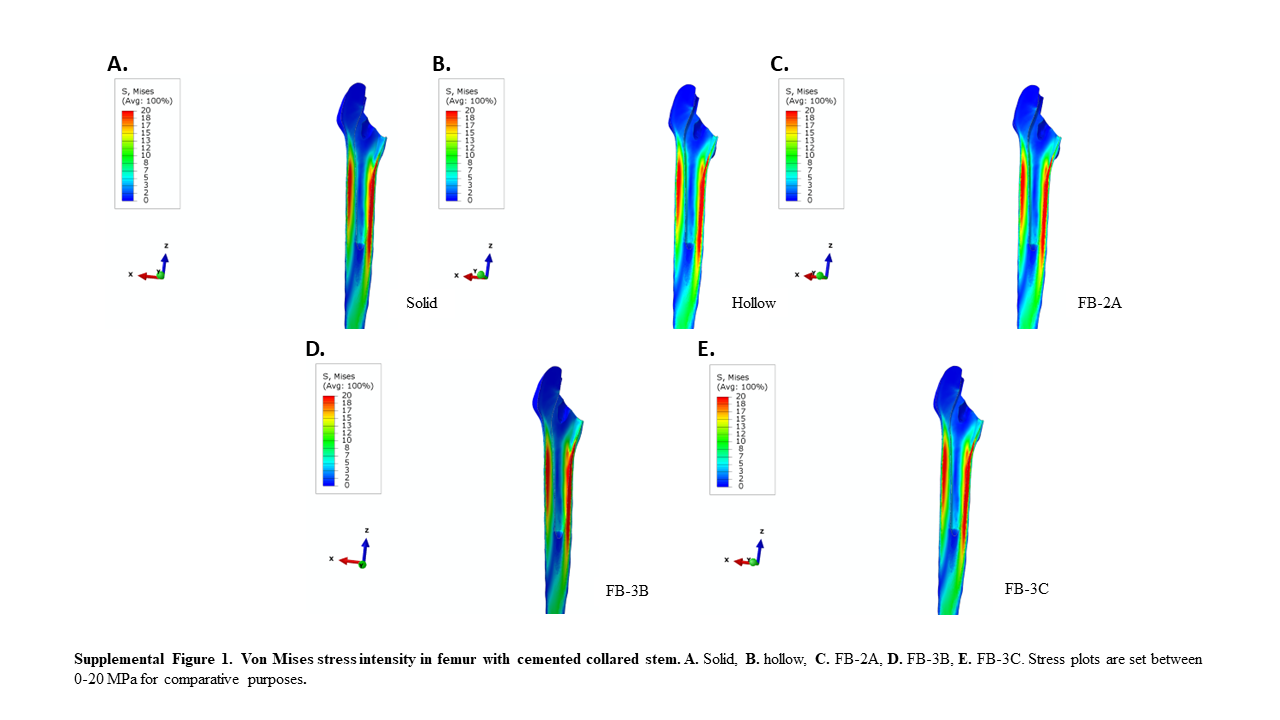

### Supplemental figure 2

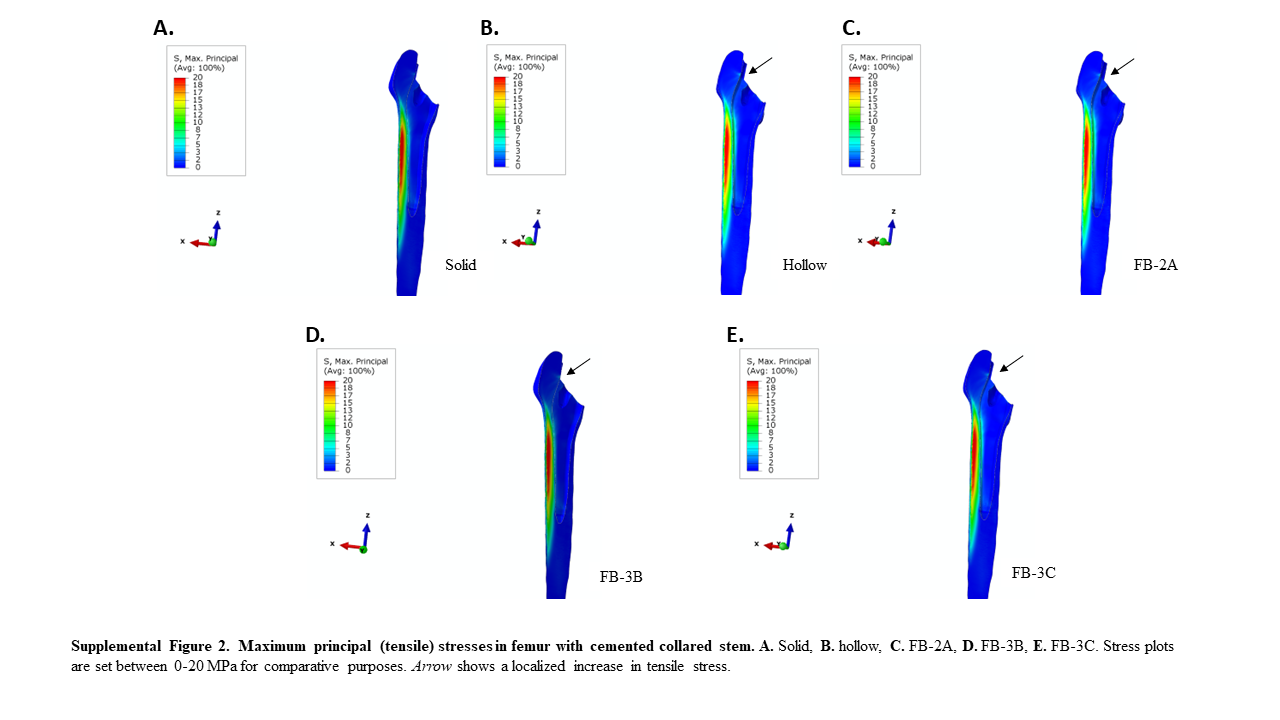

### Supplemental figure 3

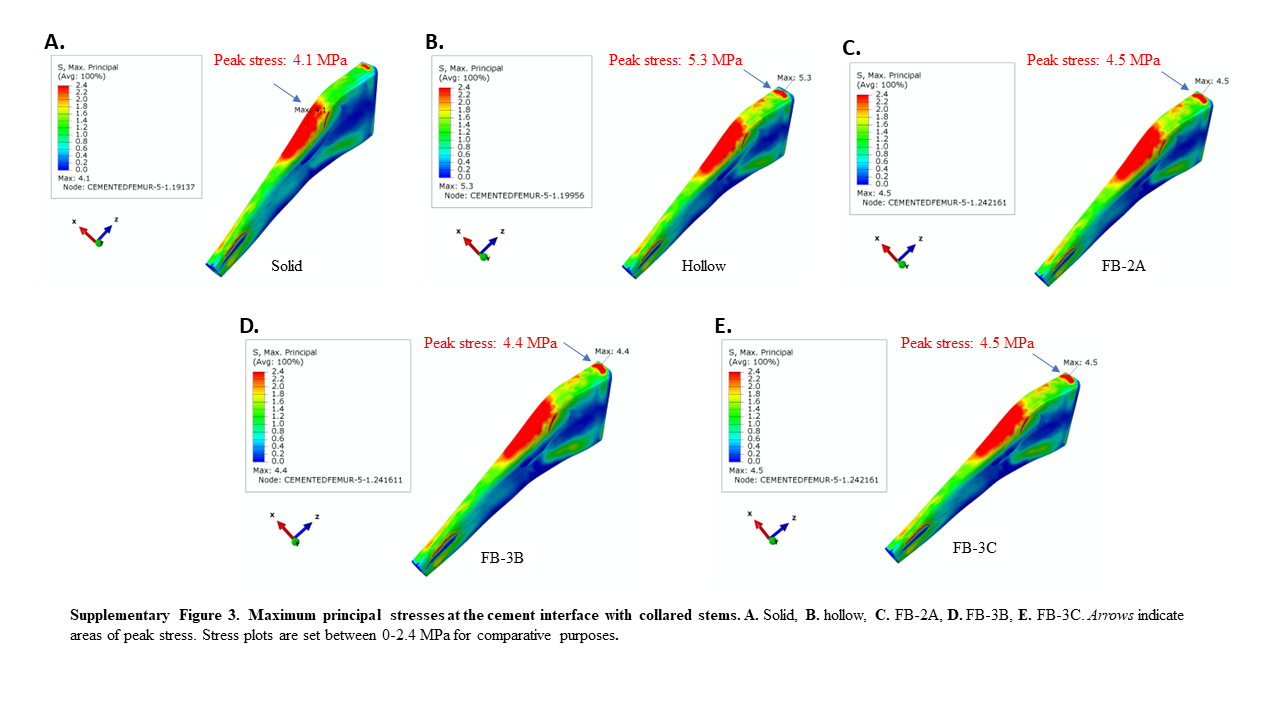

### Supplemental figure 4

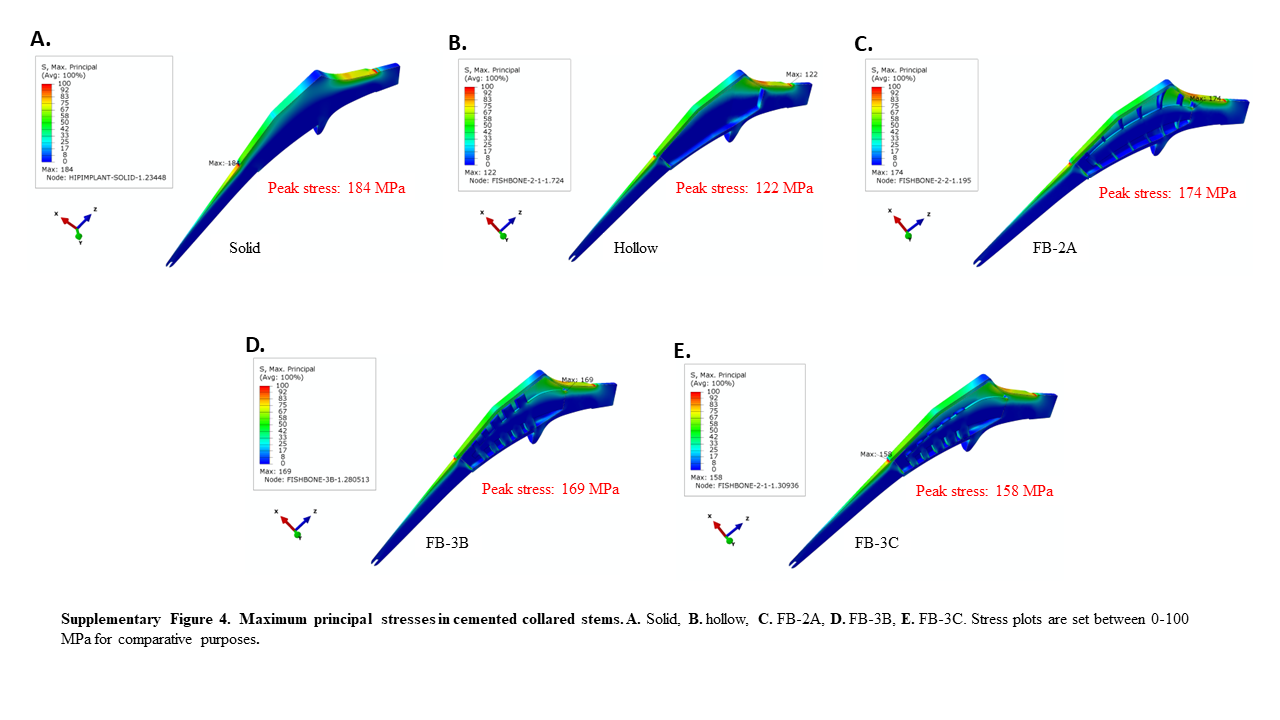

### Supplemental figure 5

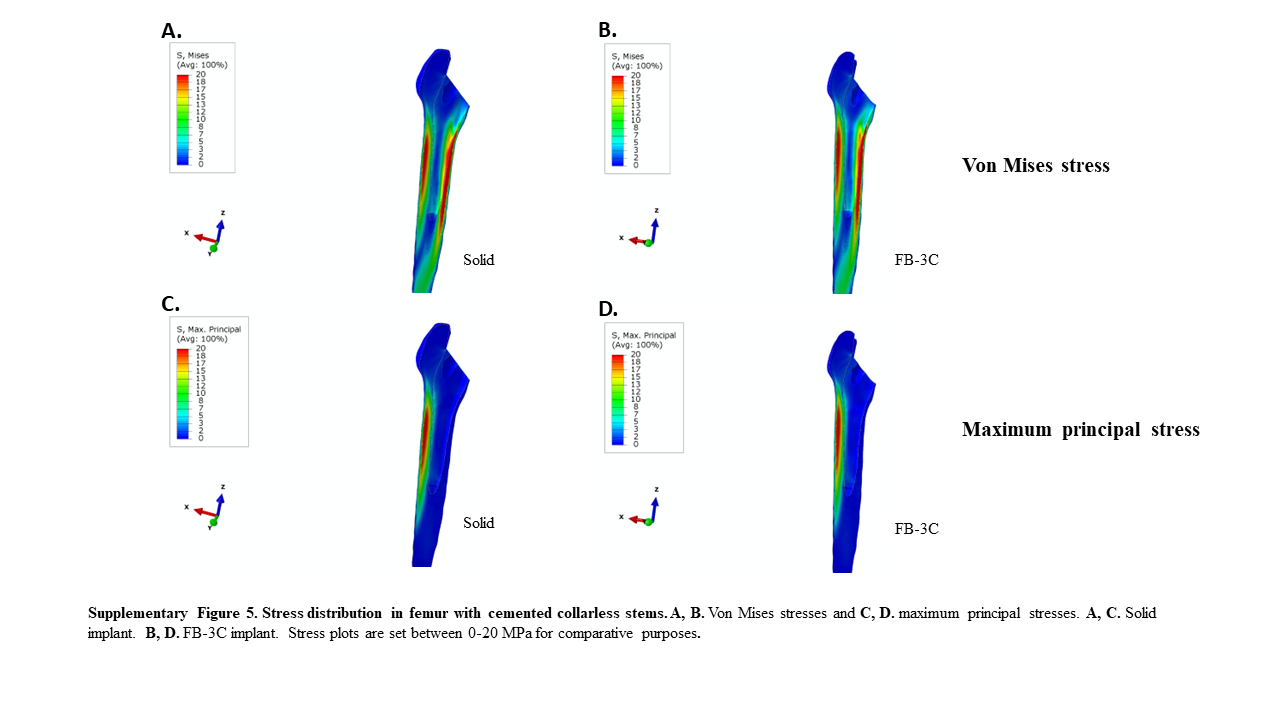

### Supplemental figure 6

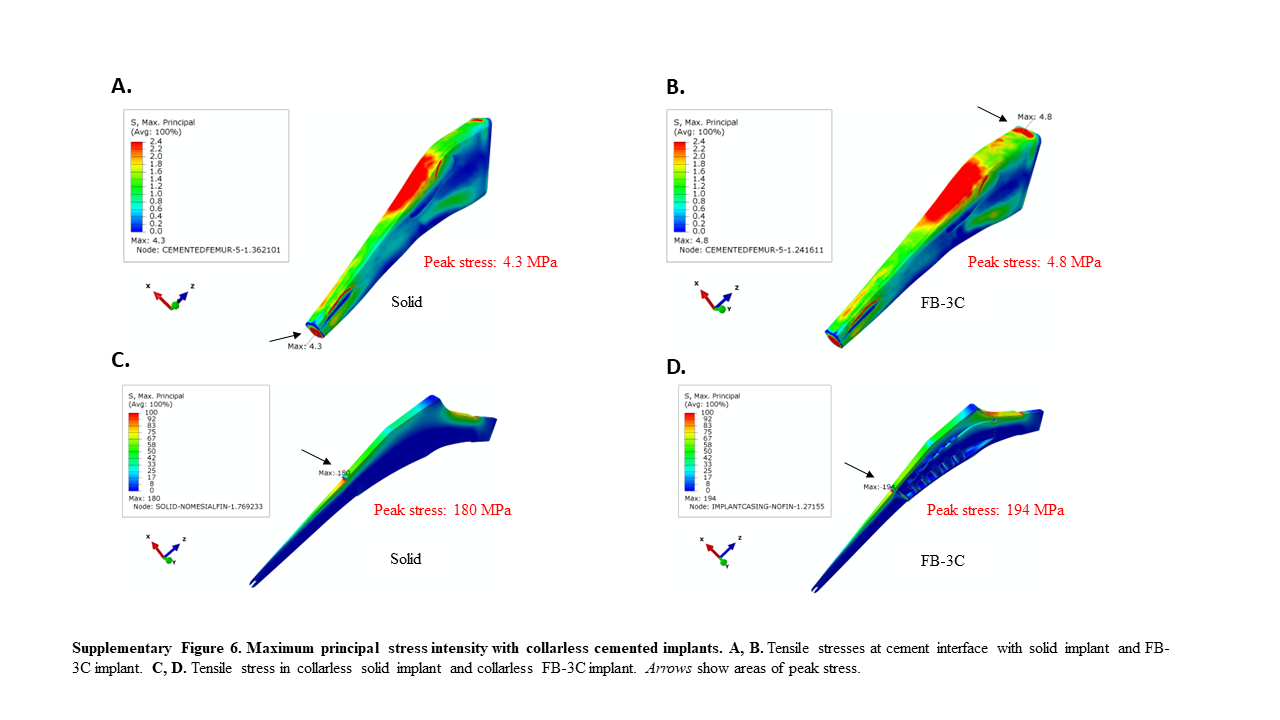
